# Task-Specialized Protein Language Models Decode the Sequence Grammar of Post-Translational Modification Sites

**DOI:** 10.64898/2026.05.08.723918

**Authors:** Subinoy Adhikari, Jagannath Mondal

**Affiliations:** Tata Institute of Fundamental Research, Hyderabad 500046, India

## Abstract

Post-translational modifications (PTMs) regulate protein signaling, localization, degradation, and cellular decision-making, yet the sequence determinants that distinguish modified from chemically eligible but unmodified residues remain difficult to decode at proteome scale. Here, we examine whether adapting a general protein language model to PTM-site prediction can reveal the biochemical logic underlying residue-level modification. We fine-tune ESM2, a protein language model trained on tens of millions of evolutionarily diverse protein sequences, for phosphorylation, acetylation, and ubiquitination-site prediction. To address the pronounced class imbalance inherent in proteome-wide PTM annotation, we combine parameter-efficient fine-tuning with focal-loss training. The resulting task-specialized models show that PTM recognition depends on model capacity, annotation depth, and modification chemistry: phosphorylation benefits from larger models, whereas acetylation and ubiquitination peak at intermediate scale. More importantly, the fine-tuned phosphorylation model exposes three layers of biological organization: it recovers canonical kinase-recognition motifs without kinase-label supervision, resolves pathway-level functional relationships among proteins from sequence-derived embeddings, and preserves evolutionary signatures of homologous phosphorylation sites across 200 eukaryotic species. These results establish task-specialized protein language models as interpretable instruments for probing PTM-site biochemistry, kinase specificity, functional organization, and evolutionary conservation.

## Introduction

Post-translational modifications (PTMs) are pivotal biochemical processes that regulate protein function and enable dynamic cellular responses to diverse physiological and environmental signals. By covalently attaching chemical groups such as phosphate, acetyl, or ubiquitin to specific amino acid residues, PTMs expand the structural and functional diversity of proteins far beyond the genomic blueprint. With over 400 distinct PTM types identified, ^1^ these modifications transform an estimated 20,000 protein-coding genes into a proteome exceeding a million unique protein species. ^2^ PTMs orchestrate critical biological processes, including signal transduction, enzymatic regulation, protein localization, and degradation, thereby serving as molecular switches that fine-tune cellular behavior.^3–5^ Their dysregulation is implicated in numerous diseases, from cancer and neurodegenerative disorders to metabolic syndromes, underscoring their significance in health and disease.^6,7^ Accurate prediction of PTM sites is essential for decoding protein function, understanding disease mechanisms, and developing precision therapeutics. Yet this prediction problem is biologically subtle: most serine, threonine, tyrosine, and lysine residues are chemically eligible for modification, but only a small subset are selected in vivo by the combined influence of local sequence context, enzyme specificity, protein function, and evolutionary constraint.

The importance of PTMs lies in their ability to introduce functional versatility without altering the genetic code. Phosphorylation regulates protein activity in pathways controlling cell division and apoptosis; ^8^ acetylation modulates chromatin structure and gene expression;^9^ and ubiquitination tags proteins for degradation, maintaining cellular homeostasis. ^10^

The combinatorial nature of PTMs, where multiple modifications coexist on a single protein, further amplifies their regulatory complexity, resembling a molecular language in which sequence context, structural environment, and modification state jointly determine regulatory meaning.^11^ This analogy is useful because PTM-site selection is also context dependent: the same residue can be modified or ignored depending on its surrounding sequence, structural environment, and regulatory role.^12^ Protein language models therefore offer a route to ask whether the biochemical grammar of PTM recognition can be inferred directly from protein sequence.

The foundation of computational PTM prediction rests upon curated databases that aggregate experimentally verified modification sites from high-throughput mass spectrometry and literature mining. General-purpose repositories such as PhosphoSitePlus, ^13^ Phospho.ELM,^14^ PHOSIDA,^15^ and the Human Protein Reference Database^16^ provide curated PTM annotations across model organisms, while Swiss-Prot/UniProtKB^17^ and RESID^18^ serve as authoritative cross-PTM knowledgebases. Modification-specific resources include UbiProt^19^ and UbiNet 2.0^20^ for ubiquitination, while integrated platforms such as SysPTM 2.0,^21^ iPTMnet,^22^ and the Eukaryotic Phosphorylation Site Database (EPSD)^23^—which alone houses over 1.6 million phosphorylation sites across 68 species—consolidate multi-PTM data with pathway and ontology annotations. Structure-resolved information is available through Phospho3D^24^ and PTM-SD,^25^ and the dbPTM resource,^26^ recently updated to dbPTM 2025,^27^ remains among the most comprehensive integrated platforms. Collectively, these databases form the training and validation backbone for downstream computational prediction tools.

Computational prediction of PTM sites has progressed from motif- and profile-based scoring to classical machine-learning classifiers and, more recently, deep neural networks that learn modification-specific sequence features directly from data. Early methods used kinase motifs, position-specific scoring, support vector machines, random forests, and physicochemical encodings to capture the local sequence environments associated with phosphorylation, acetylation, ubiquitination, and related PTMs. ^28–36^ Deep learning subsequently enabled convolutional, recurrent, and transformer-based models to learn longer-range sequence dependencies and context-dependent PTM signatures, improving residue-level prediction across multiple modification classes.^37–42^

Protein language models (PLMs) have further transformed the landscape by providing contextual residue embeddings learned from millions of natural protein sequences. These embeddings encode information about evolutionary conservation, structural constraints, domain organization, and functional context without requiring explicit supervision for each property.^43–45^ Recent PTM predictors increasingly exploit this representational capacity by combining PLM embeddings with task-specific classifier heads, multi-task learning, prompt tuning, structural features, evolutionary profiles, motif-aware modules, or unified multi-label formulations. ^46–52^ These approaches have substantially improved PTM coverage and proteome-scale applicability.

Despite this rapid progress, the current generation of PTM predictors leaves several questions open. First, the move toward broad-spectrum, multi-modal architectures has come at the cost of a clear understanding of how the underlying protein language model itself contributes to performance: when ESM2 is held fixed and combined with structural, evolutionary, or motif-aware modules, it becomes difficult to attribute gains to the language model versus to the additional modules. Second, although adapting large models for specific tasks while training only a small fraction of parameters is now well-established for natural-language models, this kind of efficient adaptation has not been systematically characterized for PTM prediction across model sizes—existing efforts largely use a single model size.^47^ Third, proteome-wide PTM prediction is intrinsically imbalanced, with phosphorylation accounting for the bulk of curated annotations, ^53^ and even within a single PTM type, the proportion of modified residues is typically below 10% of all candidate sites—a severe imbalance that standard training procedures handle poorly.^54^ Finally, most predictors are evaluated narrowly on classification metrics, with little attention to whether the learned representations recover known biological patterns such as kinase consensus motifs, functional pathway organization, or evolutionary conservation ^55^—a question that becomes especially important when the goal is not just prediction but mechanistic understanding.

Motivated by these gaps, we ask whether adapting a general protein language model to PTM-site prediction can convert it into an interpretable biological instrument for probing residue-level modification logic. We achieve this specialization through parameter-efficient fine-tuning of ESM2 family of protein language models,^44,56^ ranging from 8 million to 3 billion parameters and adapting each variant to PTM site classification using Low-Rank Adaptation (LoRA).^57^ LoRA is an efficient adaptation strategy that updates only a small fraction of model parameters and thereby makes adapting billion-parameter models practical on a single GPU. The severe imbalance between modified and unmodified residues is handled by a focal-loss training objective that gives the model stronger feedback on the rare positive cases. Within this controlled setting, we make three contributions. First, we characterize how predictive performance scales across phosphorylation, acetylation, and ubiquitination, finding that the optimal model size depends on the PTM: phosphorylation continues to improve through the 3B-parameter model, while acetylation and ubiquitination peak at 650M parameters and decline at the largest scale—a finding directly relevant to anyone choosing a model size for new PTM tasks. Second, we show that the internal patterns of attention learned by the adapted phosphorylation model recover canonical kinase consensus motifs, with different parts of the model specializing in different kinase families—providing direct mechanistic interpretability that is hard to achieve when the language model is held fixed. Third, we validate the adapted representations along two independent biological axes: similarity-based protein networks recover Gene Ontology functional organization that complements (rather than reproduces) the experimentally curated STRING interactome, and similarity at homologous phosphorylation sites correlates strongly with phylogenetic distance across 200 eukaryotic species (Figure 1). Together, these analyses establish that an adapted protein language model is valuable not only as a predictor but as an instrument whose internal representations can be read out as testable biological hypotheses—a complementary axis of value to the coverage-oriented predictors that currently define the field.

**Figure 1:**
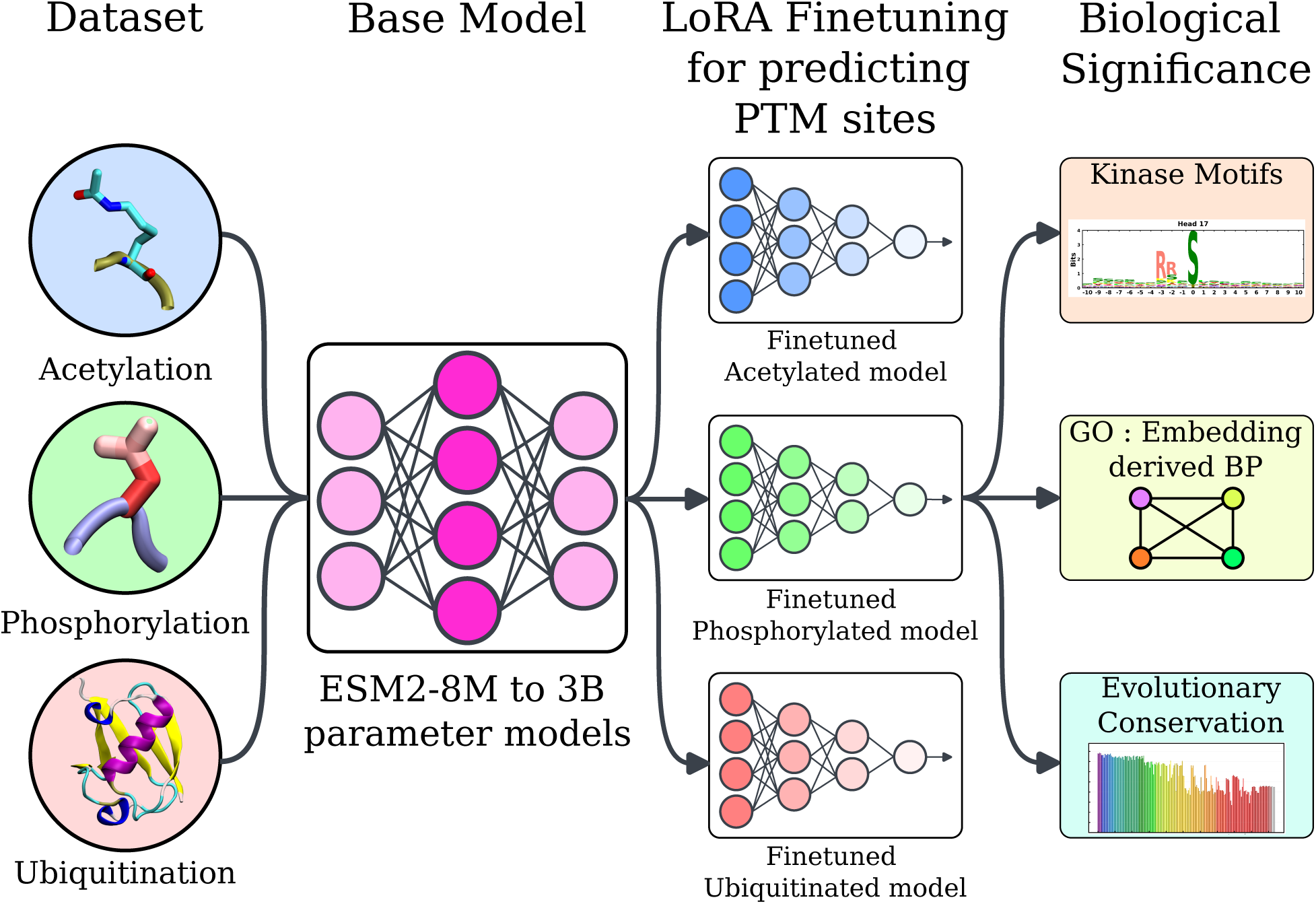
A schematic overview illustrating the development of post-translational modification (PTM)-specific predictive models. ESM2 models, ranging from 8M to 3B parameters, are LoRA-fine-tuned on datasets corresponding to three major PTMs—acetylation, phosphorylation, and ubiquitination. The resulting specialized models are used for accurate prediction of PTM sites. Downstream analyses interrogate the learned representations for three biological readouts: kinase-recognition motifs, pathway-level functional organization, and evolutionary conservation across eukaryotes.

## Results and Discussion

### Task Specialization of Protein Language Models for Residue-Level PTM Recognition

Each PTM targets a characteristic set of amino acid residues. Phosphorylation targets serine (S), threonine (T), and tyrosine (Y), wherein kinases catalyze the addition of a phosphate group to regulate protein activity and signaling. Acetylation predominantly modifies lysine (K), most prominently in histone proteins, where acetyltransferases add acetyl groups to alter chromatin accessibility and gene expression. Ubiquitination also occurs on lysine, attaching ubiquitin to mark proteins for degradation or to alter their cellular localization. Given a protein sequence, our task is therefore to ask, for every S, T, Y, or K residue, whether it is modified or not. The biological challenge is that only a small fraction of these candidate residues are actually modified under physiological conditions; the model must learn to distinguish true modification sites from the much larger pool of chemically eligible but unmodified residues based on the local sequence context surrounding each site.

We approach this problem using ESM2, a protein language model trained on tens of millions of evolutionarily diverse protein sequences. During its pre-training, ESM2 learns to predict amino acids that have been hidden from each sequence—a procedure that forces the model to internalize the recurring patterns by which residues co-occur in nature. As a result, the model’s internal representation of each residue captures information about its biochemical environment, including features related to secondary structure, residue co-evolution, and functional domains, without the model having been explicitly taught any of these concepts. The ESM2 family spans models from 8 million to 3 billion parameters (Figure 2 (a)), with larger variants showing increasingly fine-grained awareness of protein structure—a property that is useful for tasks such as PTM prediction, where the local biochemical environment is often the decisive cue.

**Figure 2:**
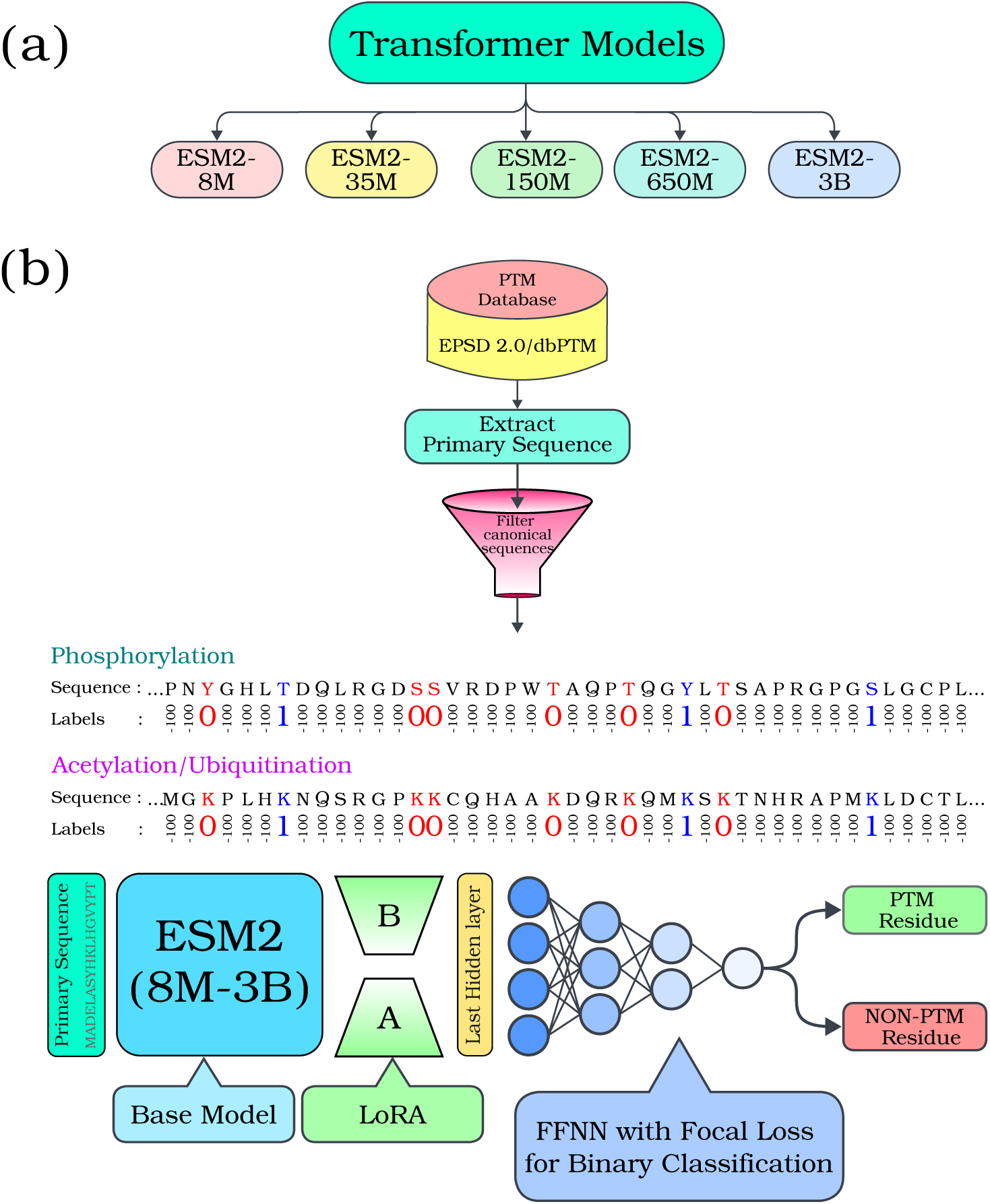
Pipeline for PTM site prediction. (a) The ESM2 family of protein language models, ranging from 8M to 3B parameters, serves as the starting point. (b) PTM datasets (EPSD 2.0 for phosphorylation; dbPTM for acetylation and ubiquitination) are processed by extracting primary sequences, filtering for canonical entries, and assigning per-residue labels (modified vs. unmodified) for each candidate residue. Each sequence is passed through the ESM2 model, whose internal representation of each residue is then used as input to a small feed-forward classifier that predicts whether the residue is modified. The ESM2 backbone is fine-tuned using LoRA, which adds a small number of trainable parameters to the model’s attention layers without altering the original weights. A focal-loss training objective addresses the pronounced class imbalance arising from the rarity of true modified residues among chemically eligible candidate sites.

A pre-trained ESM2 model is, however, a general-purpose protein representation; it captures broad sequence biology but does not by default emphasize the specific features—kinase-binding motifs, local electrostatics, sorting signals—that distinguish a true modification site.

Adapting the model to PTM prediction therefore requires further training on labeled examples. Doing so by retraining the entire 3-billion-parameter network is computationally prohibitive, while training only a small classifier on frozen ESM2 features can fail to capture the deeper sequence patterns specific to PTMs. We use a middle path: Low-Rank Adaptation (LoRA), which inserts a small set of additional trainable parameters into the model’s attention machinery while leaving the original weights untouched. This adjusts how the model attends to neighboring residues without disturbing its general protein knowledge, and reduces the training cost enough that even ESM2-3B can be adapted on a single GPU. A small classifier on top of the adapted model then produces a per-residue prediction (Figure 2 (b)). We therefore treat fine-tuning as a process of task-specialization: the model retains its broad protein-sequence knowledge but is redirected toward the residue-level biophysical features that distinguish modified from unmodified candidate sites. This proteome-wide class imbalance,where true modified residues constitute only a small minority of chemically eligible candidate sites is handled with focal loss, which gives greater weight to rare and difficult positive examples during training. Architectural and hyperparameter details are given in Methods.

### Model Capacity, Annotation Depth, and PTM Chemistry Jointly Shape Prediction Performance

The fine-tuned ESM2 models reveal a striking pattern: the benefit of fine-tuning depends strongly on PTM chemistry and data depth. In particular, model size interacts differently with each PTM, and increased capacity does not uniformly translate into improved performance. We evaluate predictions using four standard metrics, each with a clear biological interpretation. Recall measures what fraction of true modification sites the model successfully identifies—the higher the Recall, the fewer biologically real sites are missed. Precision measures what fraction of the model’s predictions are correct, capturing the rate of false positives that would mislead downstream experiments. F1-score is their harmonic mean.

The Matthews Correlation Coefficient (MCC) is a single, balanced summary statistic that performs well when modified residues are heavily outnumbered by unmodified ones, as is the case here. We report each metric averaged across both modified and unmodified residues to ensure that performance on the rare modified class is not masked by the much larger unmodified class. As a baseline, we compare against PTMGPT2,^46^ the most directly comparable transformer-based PTM predictor.

We first evaluated the performance of pre-trained baseline models on the phosphorylation site prediction task using the ESM2 models and compared them with PTMGPT2 model. As shown in Figure S1, all baseline ESM2 models (labelled as *B-ESM2* in figure S1) demonstrated poor predictive performance across all metrics. The ESM2 models achieved F1-scores ranging from 0.39 to 0.49, with AUROC values between 0.44 and 0.54, barely exceeding random chance. Precision and recall metrics were similarly underwhelming, with precision hovering around 0.48-0.61 and recall between 0.44-0.55. Most concerning were the MCC values, which ranged from −0.064 to 0.047, indicating minimal to no correlation between predictions and true phosphorylation sites. Notably, scaling to larger model sizes (650M and 3B parameters) did not yield substantial improvements, with the 650M model actually showing the worst performance (MCC: −0.064, F1: 0.44, AUROC: 0.44). In contrast, PTMGPT2 demonstrated markedly superior performance with an F1-score of 0.59, AUROC of 0.61, precision of 0.61, and MCC of 0.18, substantially outperforming all baseline ESM2 variants. However, even this best-performing baseline model exhibited limited discriminative power for phosphorylation site identification, with an MCC of only 0.182 suggesting weak predictive capability. These results indicate that general-purpose protein language models, despite being pre-trained on large-scale protein sequence data, lack the specific representations necessary for accurate phosphorylation site prediction. The poor zero-shot performance across all baseline models, particularly the near-random AUROC scores for ESM2 variants motivated our decision to fine-tune these models.

Fine-tuning the ESM2 variants on the phosphorylation dataset produced substantial improvements across all evaluation metrics and model sizes. Performance climbs from finetuned ESM2-8M through ESM2-150M (referred as *FT-ESM2*), with progressive gains in Recall and MCC indicating better sensitivity to true sites, and ESM2-3B delivers the strongest results overall (MCC 0.35, F1 0.66, highest AUROC; Figure 3 (a)). This is biologically reasonable: phosphorylation is by far the best-annotated PTM, with millions of validated sites across hundreds of kinase families, so larger models have enough labeled examples to learn the diverse sequence contexts that different kinases recognize.

**Figure 3:**
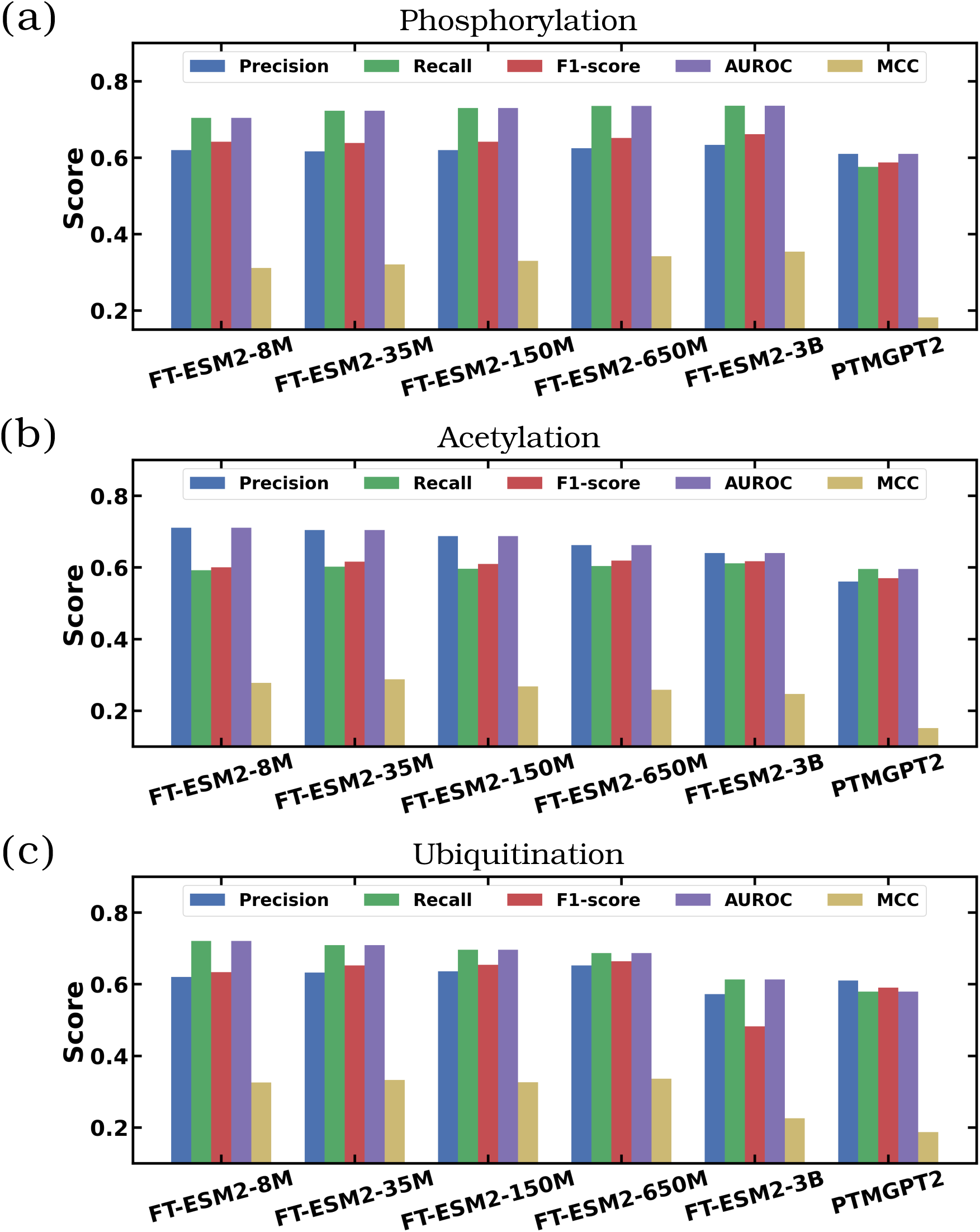
Comparison of the classification metrics for different fine-tuned ESM2 models (labelled as *FT-ESM2*) for (a) Phosphorylation, (b) Acetylation and (c) Ubiquitination. Also shown the comparison with baseline model PTMGPT2 model. ^46^

Acetylation tells a different story. Performance peaks at intermediate scale rather than continuing to improve. The smallest model behaves conservatively, predicting few sites but predicting them confidently (high Precision, lower Recall). Through ESM2-35M and ESM2-150M, F1 improves modestly while Recall remains stable; the best balance between sensitivity and specificity emerges at ESM2-650M (MCC 0.26). At ESM2-3B, all metrics drop including AUROC, suggesting that the largest model has begun to overfit to the comparatively limited acetylation training set (Figure 3 (b)).

Ubiquitination follows a similar trajectory but with a sharper drop at the largest scale. Performance improves through ESM2-150M, peaks at ESM2-650M (MCC 0.34, F1 0.66), and then degrades substantially at ESM2-3B (Figure 3 (c)). As with acetylation, this likely reflects the smaller and biologically narrower set of validated ubiquitination sites available for training—ubiquitination is harder to capture by mass spectrometry than phosphorylation, leading to less complete annotation databases.

The PTMGPT2 baseline consistently underperforms across all three PTM types when compared with fine-tuned models, with notably lower Recall indicating limited sensitivity to true modification sites and lower MCC reflecting weaker overall classification robustness compared to the LoRA-fine-tuned ESM2 models.

More broadly, these results show that the relationship between model scale and PTM prediction performance is task-dependent. Phosphorylation, with the largest and most diverse training set, benefits monotonically from increased capacity. Acetylation and ubiquitination, in contrast, peak at ESM2-650M and degrade at 3B—a pattern most consistent with overfitting in the larger model when labeled data are limited. Thus, model scaling in PTM prediction is not governed by capacity alone, but by the interplay between modification chemistry, annotation depth, and the diversity of biochemical recognition contexts represented in the training data.

### Fine-tuned Attention Heads Recover Kinase-Specific Phosphorylation Motifs

Having shown that fine-tuning improves phosphorylation prediction, we next ask whether the adapted model has learned the biochemical rules of kinase recognition rather than dataset-specific correlations. A predictor can achieve good metrics for the wrong reasons— memorizing dataset artifacts, exploiting compositional biases, or over-relying on sequence neighborhoods that happen to correlate with annotation. If the fine-tuned, task-specialized phosphorylation model has internalized biologically meaningful determinants of phosphorylation, its attention patterns should recover kinase-recognition motifs without kinase-label supervision. To distinguish biochemical learning from such shortcuts, we interrogated the model, Äôs attention patterns around annotated phosphorylation sites.

Phosphorylation is carried out by kinases, and each kinase family recognizes a characteristic short sequence motif around the phosphorylated residue—for example, the proline-directed sites preferred by MAPKs and CDKs, or the basic residues upstream of PKA and PKC sites. These motifs were never given to the model during training; the labels indicate only whether a site is phosphorylated. If the adapted model has internalized real biology, the way it weights the residues surrounding a phosphorylation site—through the model’s internal “attention” mechanism, which encodes how much each residue’s representation depends on each other residue in the sequence—should reproduce these consensus motifs without any supervision over kinase identity. We therefore aggregated these attention values from the final layer of the adapted ESM2-3B model across all annotated phosphorylation sites, within a ±10-residue window, and converted the resulting positional preferences into information-content sequence logos. The model has 40 such attention patterns running in parallel (the “heads”), and each head can be inspected separately to see what it has learned to focus on (Figure 4 (a); details in Methods).

**Figure 4:**
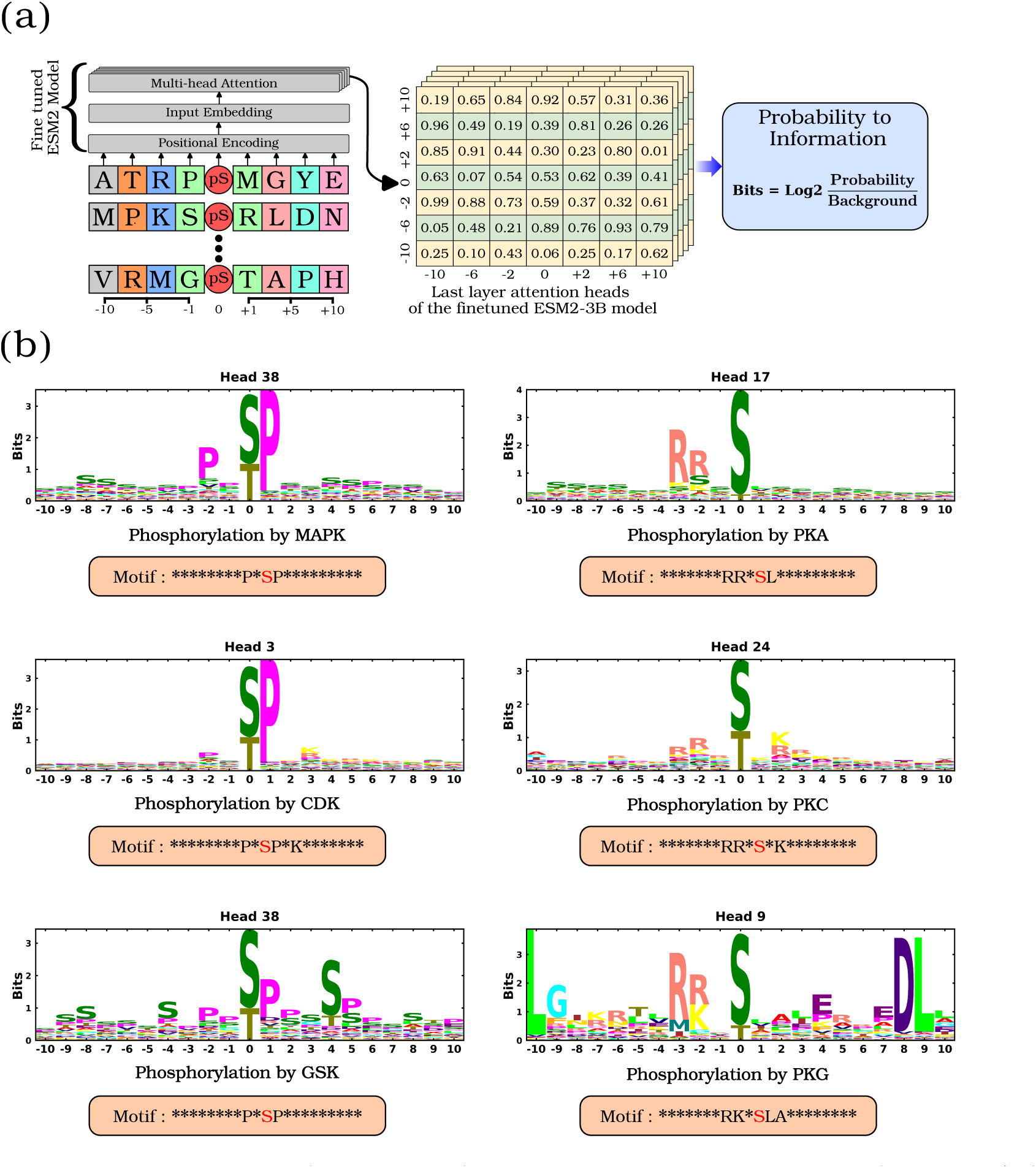
Kinase consensus motifs recovered from internal attention patterns of the LoRA-fine-tuned ESM2 model. (a) Schematic of the analysis: protein sequences containing phosphorylated residues are passed through the fine-tuned model, and attention values from each phosphorylation site to its surrounding residues (−10 to +10 positions) are extracted from the model’s final layer (40 attention heads). These values are aggregated across all annotated sites in the dataset to produce position-specific amino acid preferences, which are then converted into information-content sequence logos. (b) Sequence logos derived from these patterns reveal the canonical kinase consensus motifs (e.g. MAPK, CDK, GSK, PKA, PKC, PKG) that the model has implicitly learned to associate with phosphorylation, despite never having been told which kinase modifies which site.

Across the major kinase families known to leave distinct sequence signatures, the resulting logos cleanly recover the canonical motifs (Figure 4 (b)). The CMGC family—which includes CDKs,^54^ MAPKs,^54^ and GSKs^58^—is well known for proline-directed phosphorylation, and the model recovers this through a single dedicated attention head that captures the canonical P*SP* motif (proline at −2 and +1) for both MAPK and GSK substrates, while a separate head captures the CDK-specific extension P*SP*K. The AGC family—PKA,^59^ PKG,^59^ and PKC^60^—is characterized by basic residues upstream of the phosphorylation site, and each member is captured by a distinct head with a refined version of this pattern (RR*SL for PKA, RK*SLA for PKG, RR*S*K for PKC). The CAMK family ^61^ similarly relies on an upstream basic residue, recovered as the R**S motif for both CAMKL and CAMK2.

Beyond these family-level patterns, individual kinases also receive dedicated representations: DMPK,^62^ MAPKAPK,^63^ and AKT each show characteristic basic motifs, and casein kinase 2 (CK2) recovers its acidic SD*E pattern (Figure S2). The one notable exception is casein kinase 1 (CK1), for which the model did not recover a clear consensus, consistent with CK1, Äôs known reliance on prior phosphorylation at, àí3 rather than a fixed primary-sequence motif.

Two points are worth highlighting. The model recovers known kinase motifs without ever being told which kinase modifies which site, and different parts of the model (different heads) specialize in different kinase signatures, suggesting that the model learns a divided-up representation of phosphorylation determinants rather than a single universal motif. Together, these observations show that fine-tuning exposes kinase-specific biochemical specificity within the model: distinct attention heads recover distinct phosphorylation-recognition motifs without kinase-label supervision. Although attention weights do not provide a complete mechanistic explanation of model decisions, their recovery of known kinase consensus motifs provides a biologically grounded readout of the sequence features emphasized by the fine-tuned model.

### Task-Specialized Embeddings Resolve Pathway-Level Functional Organization

The previous section showed that the adapted model captures kinase-specific motifs at the residue level. We next asked whether task-specialization also reorganizes the protein-level embedding space in a biologically meaningful way. In particular, we therefore asked whether the same fine-tuned representation also encodes higher-order functional organization: specifically, whether proteins annotated to related biological processes occupy proximal regions of the model-derived embedding space.

To test this, we represented each protein as a single embedding vector and constructed a graph in which edges connect pairs of proteins with sufficiently similar embeddings. If the fine-tuned representations encode biologically meaningful functional information, proteins participating in the same biological process should be preferentially connected within this graph. This analysis requires both functional annotations and a reference network for comparison. For functional labels, we used the Gene Ontology (GO) Biological Process namespace,^64^ a hierarchical vocabulary in which proteins are annotated according to the biological processes in which they participate. As a reference network, we used the STRING protein–protein interaction database, ^64^ restricted to high-confidence interactions, which captures physical interactions as well as curated functional associations.

Our comparison rests on three metrics from the framework of Chagoyen and Pazos. ^64^ For a given GO term (e.g. “cell cycle”), we extract the subgraph induced by proteins annotated to that term and compute: (i) the *density* (*D*), defined as the fraction of possible edges among those proteins that are present; (ii) the *clustering coefficient* (*CC*), which quantifies the extent to which neighbors of a protein are themselves connected, thereby measuring local triangle-richness; and (iii) the *segregation ratio* (*S_r_*), which measures whether a protein’s connections are preferentially concentrated within its annotated GO term relative to chance expectation. Thus, density and clustering coefficient quantify local cohesiveness within a functional category, whereas segregation ratio quantifies the specificity with which that category is separated from the rest of the network.

To build the embedding-derived network, we represented each protein by averaging the residue-level representations from the adapted ESM2 3B model into a single vector per protein and connected pairs of proteins whose vectors exceeded a calibrated cosine-similarity threshold. The threshold was chosen so that the embedding-derived network and the STRING network contained comparable numbers of edges over the same set of proteins. This calibration controls for trivial differences in global connectivity and allows the comparison to focus on how each network organizes proteins with respect to biological function (Figure 5 (a); details in Methods).

**Figure 5:**
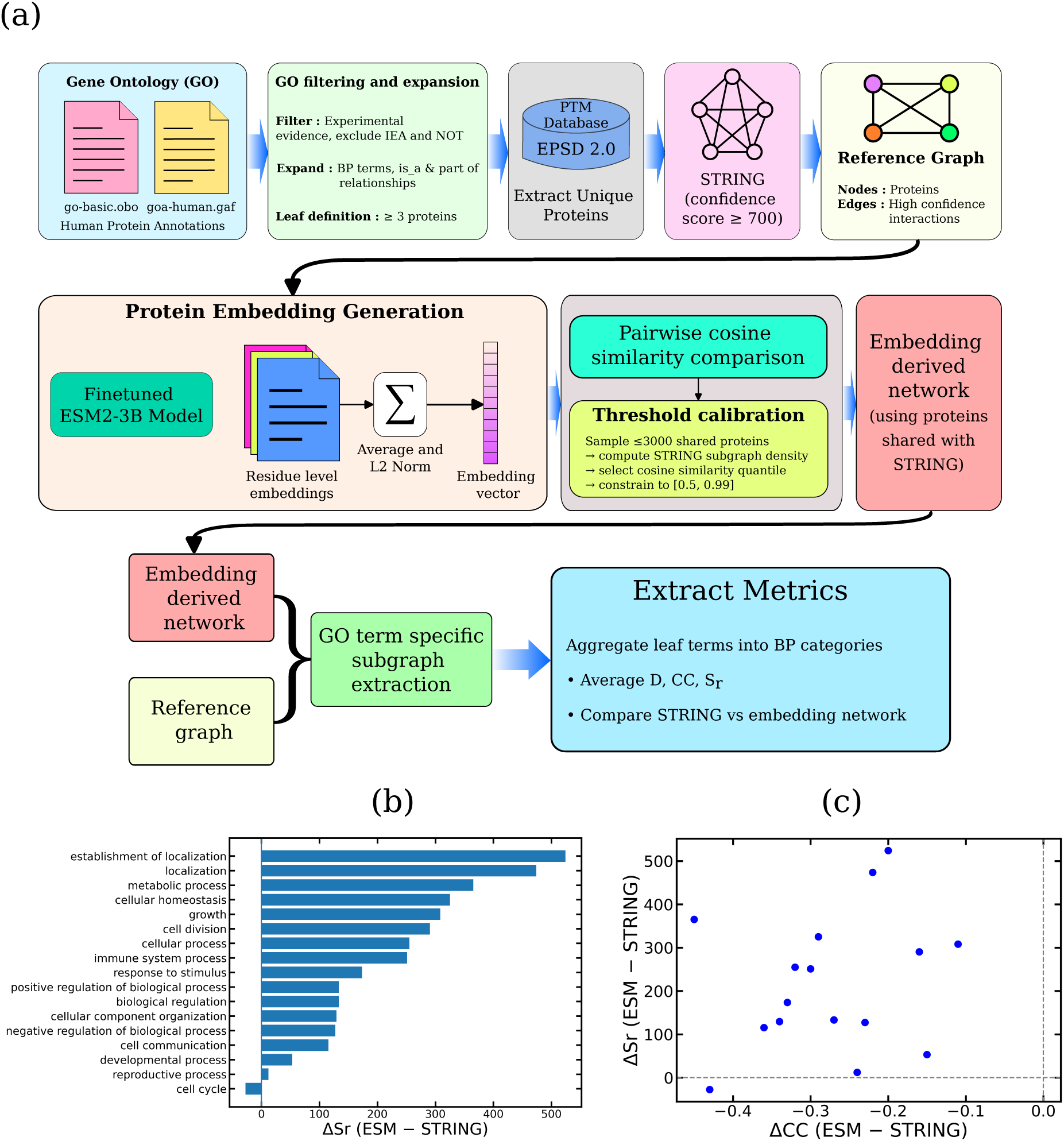
Methodological workflow and comparative analysis of functional coherence. (a) Protein function annotations were taken from the Gene Ontology (go-basic.obo, goa_human.gaf), restricted to terms supported by experimental evidence and to proteins present in EPSD 2.0. A high-confidence STRING interaction network (score ≥ 700) was constructed alongside a similarity network built from the adapted ESM2 3B representations. The similarity threshold for the second network was calibrated to match the STRING network’s edge density on the shared protein set, so that the two networks are directly comparable. Functional coherence within each Gene Ontology biological process category was quantified using three metrics: density (*D*), clustering coefficient (*CC*), and segregation ratio (*S_r_*). (b)–(c) Differences (Δ) between the similarity network and STRING for representative biological process categories. The trend toward Δ*S_r_ >* 0 and Δ*CC <* 0 indicates that the similarity network is more functionally specific (proteins in the same category are more selectively connected) but locally sparser (fewer triangles among them) than STRING.

The two networks reveal distinct but complementary modes of biological organization (Figure 5 (b)–(c)). Across nearly every biological process category, the network derived from model representations shows substantially higher segregation than STRING, indicating that proteins annotated to the same biological process are more selectively connected in the fine-tuned embedding space. The largest gains are observed for transport- and localization-related categories, including *establishment of localization* and *localization*, with several-fold increases. Broader categories such as *immune system process* and *cellular homeostasis* also show large gains (Table S1). These trends suggest that the model’s sequence representations encode determinants of biological function such as signal peptides, transmembrane domains, sorting motifs, conserved family signatures, and other sequence-level features that may define functional categories without necessarily producing direct physical interactions.

The increase in functional segregation is accompanied by a systematic reduction in local cohesiveness, with average decreases of roughly 0.27 in clustering coefficient and 0.21 in density across categories. This behavior is consistent with the different information content of the two networks. STRING integrates physical interactions, curated pathway memberships, and experimental evidence, and therefore tends to form dense, triangle-rich modules corresponding to protein complexes and pathway neighborhoods. In contrast, the embedding-derived network connects proteins on the basis of similarity in the fine-tuned sequence representation space. Such edges can reflect shared functional or evolutionary signatures, but they need not correspond to direct interaction partners or closed local interaction neighborhoods. The *metabolic process* category illustrates this distinction: STRING places metabolic proteins in dense pathway-like modules, whereas the embedding-derived network links them more sparsely while still preserving the overall functional grouping.

The *cell cycle* category provides an informative boundary case, as it is the only major category in which the segregation ratio decreases in the embedding-derived network relative to STRING. This result is consistent with the biological nature of cell-cycle regulation, which depends heavily on transient assemblies, cyclin–CDK partnerships, checkpoint complexes, and context-dependent interaction states. Such organization is more directly captured by experimental interaction networks than by primary-sequence-derived representations alone. Thus, rather than indicating a general failure of the embedding-derived network, the cell-cycle case delineates the boundary between functional similarity encoded in sequence and cellular organization that depends on dynamic interaction architecture.

Taken together, the embedding-derived network captures a complementary axis of biological organization—one that emphasizes shared sequence, evolutionary, and functional signatures over physical interaction architecture. Rather than reproducing curated interaction databases, it provides an alternative representation in which functional groupings emerge directly from the fine-tuned protein language model embedding space. Thus, the fine-tuned representation does not simply reproduce curated interaction databases. Instead, it reveals a sequence-encoded layer of pathway organization that is most apparent for processes shaped by shared localization signals, family signatures, and regulatory sequence features.

### Adapted Representations Preserve Evolutionary Constraints on Phosphorylation Sites

A natural concern about any deep learning model trained on a fixed dataset is whether it has learned biology or whether it has merely memorized statistical patterns specific to the training distribution. The previous two analyses have already shown that the fine-tuned model recovers known kinase motifs and meaningful functional groupings. Here we apply a third, independent test: does the model encode features that are conserved across evolution? The logic of the test is straightforward. Phosphorylation is an ancient regulatory mechanism, and many phosphorylation sites are functionally conserved across distantly related eukaryotes even when the surrounding sequence has diverged. If task-specialization has captured the biophysical determinants of phosphorylation rather than memorizing human-specific patterns, homologous phosphorylation sites should remain proximal in the fine-tuned embedding space in a manner that decays with phylogenetic distance. If, on the other hand, the model has merely memorized human-specific patterns, similarity should drop off sharply as soon as we leave the training distribution, with no smooth dependence on phylogeny.

To test this, we assembled a panel of 200 eukaryotic species spanning 41 major taxonomic groups—vertebrates, invertebrates, fungi, protists, and plants—representing over a billion years of evolutionary divergence (Figure 6). For each human phosphorylation site in a curated, non-redundant set, we identified the homologous residue in each organism via BLASTP and multiple sequence alignment. We then extracted the fine-tuned model’s internal representation (a numerical vector) for both the human residue and its organism counterpart, and measured how similar the two vectors point in their high-dimensional space (the cosine of the angle between them, with 1 meaning identical and 0 meaning unrelated). We averaged these similarities at the organism level and asked whether the resulting score correlates with phylogenetic distance from human (Figure 7, S3, S4; details in Methods).

**Figure 6:**
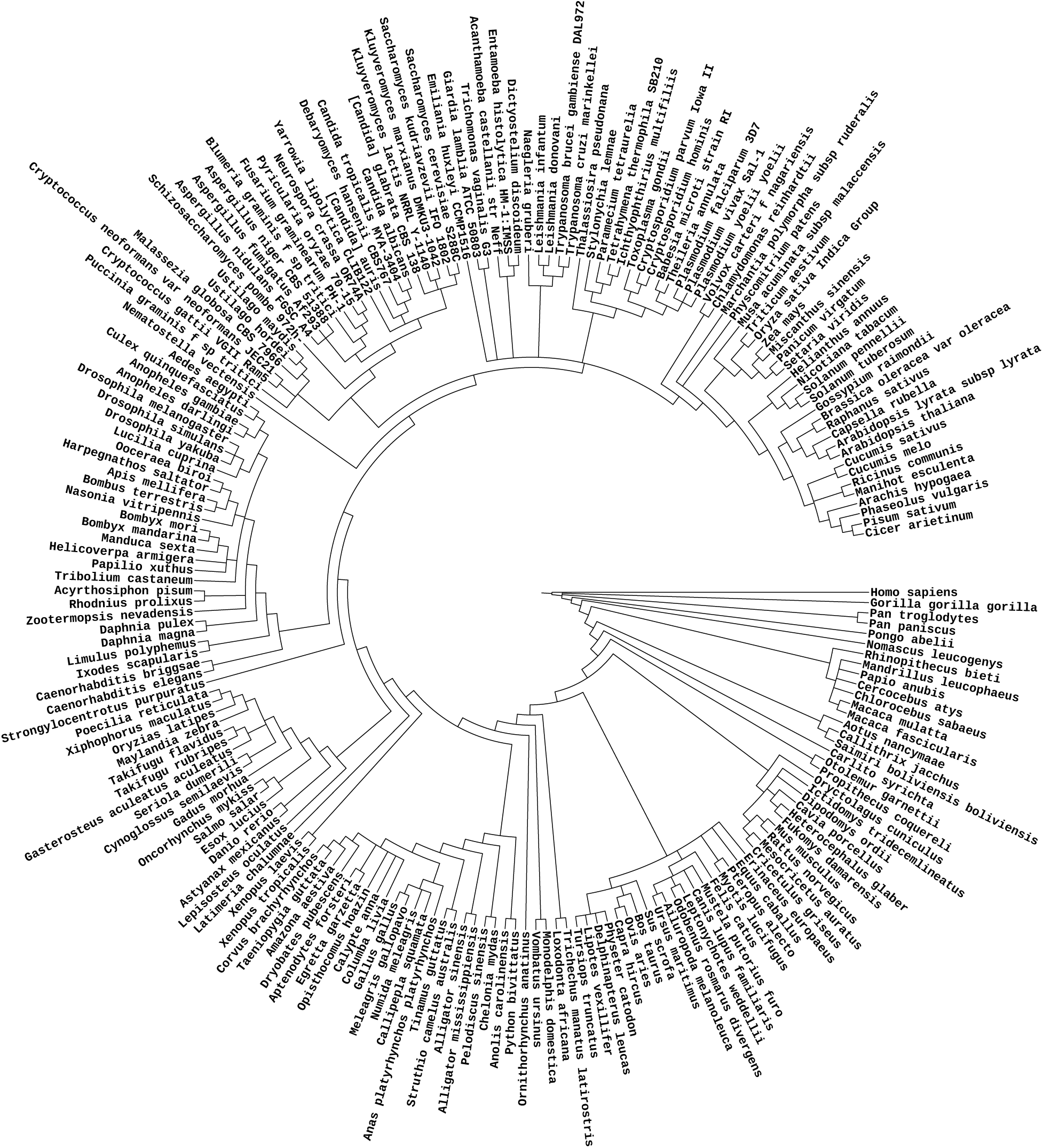
The circular phylogenetic tree illustrates the broad evolutionary relationships and genetic distances across a diverse range of eukaryotic species, with root at Homo Sapiens

**Figure 7:**
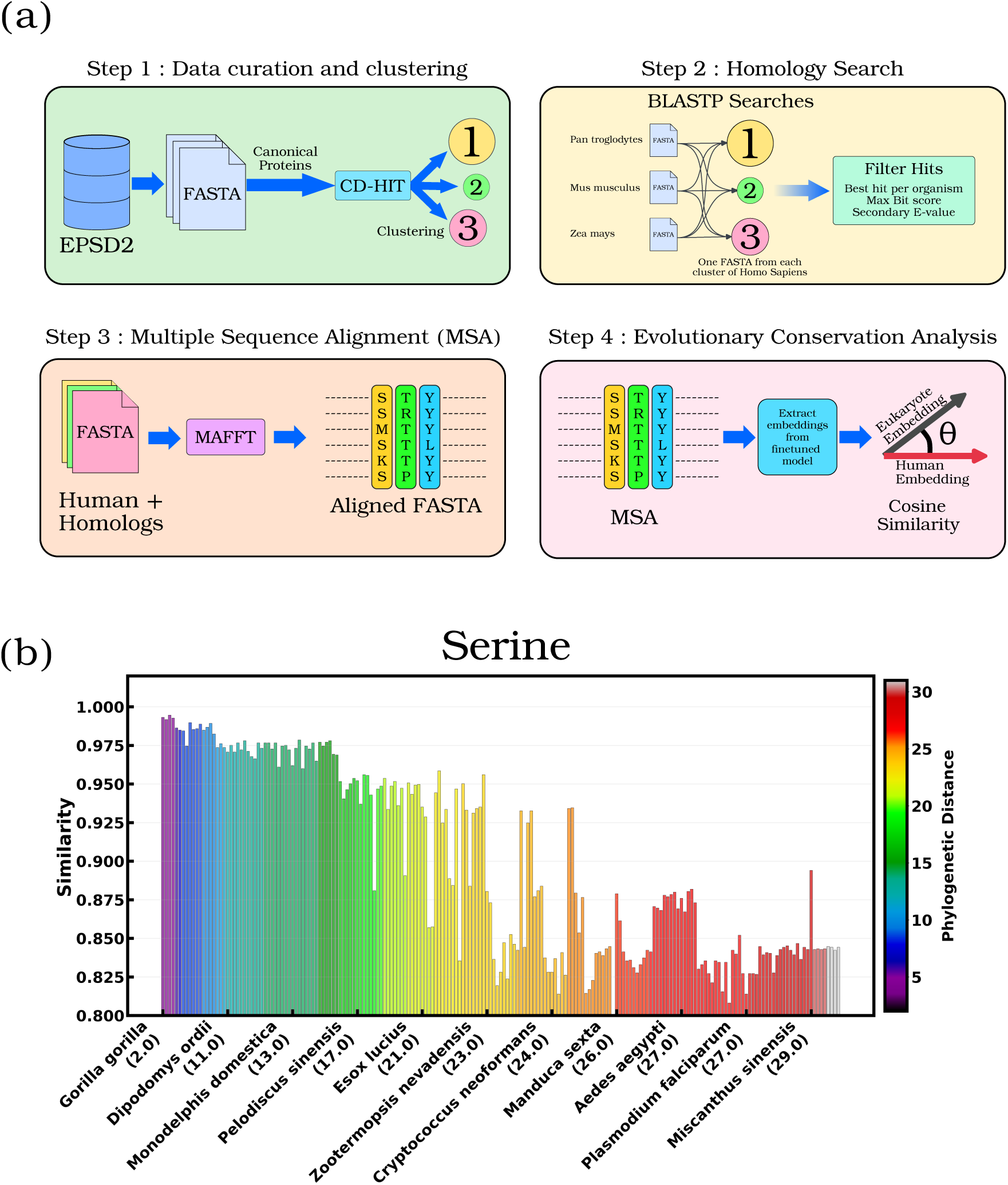
Overview of the evolutionary analysis pipeline for phosphorylation site conservation across eukaryotes. (a) Pipeline. Experimentally validated human phosphorylation sites from EPSD2 are filtered to canonical proteins and clustered with CD-HIT to remove redundancy. Representative sequences from each cluster are used to identify homologs across the panel of eukaryotic species via BLASTP, followed by selection of the best hit per organism and construction of multiple sequence alignments with MAFFT. The fine-tuned model is used to produce its internal representation for both the human residue and its homologous counterpart at each aligned phosphorylation site, and the similarity between the two is quantified by the cosine of the angle between the two representation vectors. (b) Average similarity at serine phosphorylation sites between human and homologs across the panel of eukaryotic species, with bars colored by phylogenetic distance from human and ordered along the eukaryotic tree.

This analysis reveals a strong, clean negative correlation between phylogenetic distance and similarity for all three phosphorylatable residues: serine (*r* = −0.86), threonine (*r* = −0.85), and tyrosine (*r* = −0.84). The consistency of these correlations across residue types is itself meaningful—the same evolutionary signal is recovered regardless of which amino acid gets phosphorylated. As organisms diverge from humans, the model’s representations of corresponding phosphorylation sites become smoothly less similar, mirroring the gradual accumulation of sequence and structural changes at these regulatory positions. The fact that the trend extends from closely related primates all the way to deeply divergent plant and protist lineages—rather than collapsing to noise outside the training distribution—indicates that the model has learned features of phosphorylation that predate the human lineage by hundreds of millions of years. This provides an independent evolutionary validation that the task-specialized representation captures conserved determinants of phosphorylation-site recognition rather than merely human-specific or dataset-specific patterns.

## Conclusion

We have shown that parameter-efficient fine-tuning can convert a general protein language model into an interpretable instrument for PTM-site prediction and biological discovery. By fine-tuning the ESM2 family from 8M to 3B parameters with LoRA and a focal-loss objective that handles the rarity of true PTM sites, we obtain strong residue-level predictions for phosphorylation, acetylation, and ubiquitination on a single GPU. More importantly, this design provides a clean setting in which the contribution of the PLM backbone, model scale, and PTM-specific data depth can be examined directly.

A central practical finding is that the optimal model size depends on the PTM. Phosphorylation, with the largest and most diverse training set, benefits all the way to ESM2-3B, which gives the best metrics. Acetylation and ubiquitination peak at ESM2-650M and decline at 3B, a pattern most consistent with the larger model overfitting when labeled data are limited. The practical take-away is that backbone size should be matched to the PTM and the amount of labeled data available rather than defaulting to the largest available model.

Beyond classification, three independent biological readouts show that the fine-tuned phosphorylation model learns interpretable determinants of PTM-site recognition. Inside the model, fine-tuned attention patterns recover canonical kinase consensus motifs—proline-directed for CMGC kinases, basic upstream for AGC and CAMK families, acidic for CK2— without the model ever being told which kinase modifies which site. Networks built from similarity between fine-tuned protein representations recover Gene Ontology Biological Process organization with substantially higher functional coherence than the experimentally curated STRING interactome, while remaining sparser locally; the one major exception— cell cycle, dominated by transient assemblies that primary sequence does not encode—marks a sensible boundary of what sequence-based representations can recover. Finally, similarity between human and homologous phosphorylation sites correlates strongly and consistently with phylogenetic distance (*r* ≈ −0.85) across 200 eukaryotic species, indicating that the model has learned phosphorylation features conserved across more than a billion years of evolutionary divergence.

More broadly, this study establishes fine-tuning as a route for transforming general protein language models into biological instruments. Once adapted to PTM-site recognition, their embeddings and attention patterns can be interrogated to reveal kinase specificity, pathway-level functional organization, and evolutionary conservation. Task-specialization therefore provides a framework not only for residue-level PTM prediction, but for extracting mechanistic hypotheses about protein regulatory biochemistry from sequence-trained models.

## Methods

### Dataset Curation and Handling Class Imbalance

The model is trained and evaluated on curated datasets for three major post-translational modifications (PTMs): phosphorylation, acetylation, and ubiquitination. Phosphorylation data were obtained from the Eukaryotic Phosphosite Database (EPSD 2.0),^65^ which provides experimentally validated sites across 223 eukaryotic species. Acetylation and ubiquitination data were retrieved from dbPTM,^27^ a comprehensive repository of experimentally verified PTM annotations.

#### Phosphorylation Dataset

The initial dataset comprised 362,707 proteins, which were refined to 327,516 unique sequences after resolving duplicate groups and removing redundant entries. A quality control step further excluded 48 invalid PTM annotations that did not correspond to serine (S), threonine (T), or tyrosine (Y) residues. In total, 35,760,439 candidate residues were identified, including approximately 19.1 million serines, 11.2 million threonines, and 5.3 million tyrosines. Among these, 2,573,592 (7.2%) were experimentally validated phosphorylation sites, while the remaining 33,186,847 (92.8%) were non-phosphorylated, resulting in a negative-to-positive ratio of 12.9:1. This imbalance is particularly severe for tyrosine residues (3.6% phosphorylated), compared to serine (9.3%) and threonine (5.4%).

The dataset was partitioned into training, validation, and test sets in an 80:10:10 ratio, stratified by protein sequence to prevent data leakage. The training set contains 262,012 sequences (28,584,069 residues, 2,055,505 positives, 7.19%), while the validation and test sets each contain 32,752 sequences with comparable residue distributions and positive rates (7.22%). Residue-specific proportions are preserved across splits, ensuring consistent evaluation.

#### Acetylation Dataset

The acetylation dataset consists of 35,572 protein sequences with 21,269,017 total residues, including 1,480,397 lysine (K) residues. Among these, 110,386 (7.5%) are acetylated, yielding a negative-to-positive ratio of 12.4:1. The data were split into training (28,457 sequences), validation (3,557 sequences), and test (3,558 sequences) sets using the same stratified 80:10:10 scheme. The class distribution remains consistent across splits, with approximately 7–8% of lysines labeled as modified.

#### Ubiquitination Dataset

The ubiquitination dataset includes 30,874 protein sequences comprising 17,481,793 residues and 1,084,833 lysine residues. Of these, 118,293 (10.9%) are ubiquitinated, resulting in a comparatively less severe imbalance (8.2:1). The dataset is similarly partitioned into training (24,699 sequences), validation (3,087 sequences), and test (3,088 sequences) sets, maintaining consistent class proportions across all splits.

A key challenge across all datasets is the pronounced class imbalance inherent in proteome-wide PTM prediction: true modified residues are rare relative to the much larger pool of chemically eligible but unmodified candidate sites.^66^ Standard loss functions such as cross-entropy tend to bias models toward the majority class, leading to poor sensitivity for rare but biologically significant modification sites. To address this, we employ a weighted focal loss function^67^ with focusing parameter *γ* = 2.0:

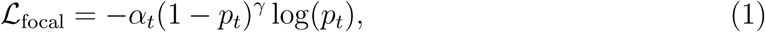

where *p_t_* denotes the predicted probability of the true class and *α_t_* is a class-specific weighting factor. Class weights are set to 1 and 12.9 for negative and positive samples in phosphorylation, 1 and 12.4 for acetylation, and 1 and 8.2 for ubiquitination, reflecting their respective imbalance ratios. Padding tokens and masked residues are excluded from loss computation to ensure that only valid positions contribute to training.

Overall, the framework combines the evolutionary and structural representations captured by ESM2^44^ with parameter-efficient fine-tuning via LoRA^57^ and imbalance-aware optimization through weighted focal loss. This design enables robust residue-level prediction performance, maintaining both precision and sensitivity in large-scale, highly imbalanced PTM datasets.

### Model Architecture and Fine-Tuning Strategy

Pre-trained ESM2 models (8M, 35M, 150M, 650M, and 3B parameters) were used as backbone encoders for contextual residue-level embeddings and fine-tuned with LoRA. In this parameter-efficient fine-tuning scheme, the original backbone weights remained frozen while trainable low-rank updates were introduced into the transformer layers.

LoRA modules were inserted into the query, key, value, and output projection layers of each attention block.^56^ Each adapter consisted of a rank-*r* decomposition with *r* = 16 and scaling factor *α* = 32. The original backbone weights were kept frozen, while LoRA parameters and all bias terms were trainable. The per-residue classification head was implemented as a feed-forward neural network (FFNN), with architecture dependent on the embedding dimension of each ESM2 variant:

- ESM2-8M / 35M: 256 → 128 → 32 → 16 → 8 → 4 → 2
- ESM2-150M: 512 → 256 → 128 → 32 → 16 → 8 → 4 → 2
- ESM2-650M: 1024 → 512 → 256 → 128 → 32 → 16 → 8 → 4 → 2
- ESM2-3B: 2048 → 1024 → 512 → 256 → 128 → 32 → 16 → 8 → 4 → 2

Each layer consisted of a linear transformation followed by layer normalization ^68^ and Tanh activation. Linear weights were initialized using Xavier uniform initialization ^69^ and biases were initialized to zero. Protein sequences were preprocessed by replacing rare amino acids (O, B, U, Z, J) with “X”. Tokenization was performed using the ESM2 tokenizer with a maximum sequence length of 1024 tokens. To ensure alignment with special tokens, residue-level labels were truncated to 1022 positions and padded with ignore indices (−100) at both termini. A custom DataCollatorForTokenClassificationESM was used to maintain consistency between input IDs, attention masks, and label tensors. To address class imbalance, a weighted focal loss was used with *γ* = 2.0. Loss computation excluded padding and masked positions (−100). Training was performed using mixed-precision (FP16) to improve computational efficiency^70^ on NVIDIA A6000 GPUs. The AdamW optimizer^71^ was used with a learning rate of 5 × 10*^−^*^5^ and weight decay of 1.0. A batch size of 2 was used without gradient accumulation. Models were trained for up to 10 epochs with early stopping^72^ based on validation loss, using a minimum improvement threshold of 0.005 and patience of 5 evaluation steps. Evaluation was conducted every 1250 training steps. Metrics were computed only on valid (non-padded) residue positions. Regularization was provided by the intrinsic dropout layers in ESM2 (dropout rate = 0.1)^73^ and layer normalization in the classifier. No additional explicit regularization was applied. All models were implemented in PyTorch ^74^ using the HuggingFace Transformers library.^75^ Custom components included the focal loss function, data collator, and early stopping mechanism. All experiments were performed with fixed random seeds for PyTorch, NumPy, and Python to ensure reproducibility. Model performance was evaluated using Precision, Recall, F1-score, and Matthews Correlation Coefficient (MCC),^76^ computed from true positives (TP), true negatives (TN), false positives (FP), and false negatives (FN):

**Positive class metrics:**

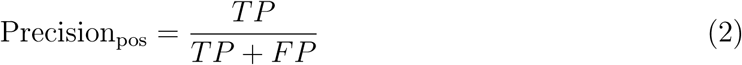

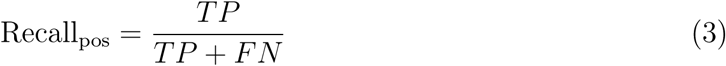

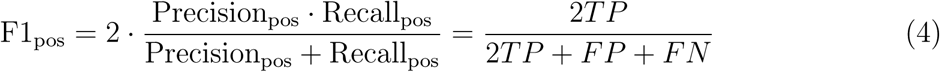

**Negative class metrics:**

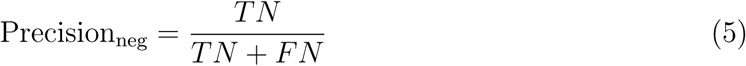

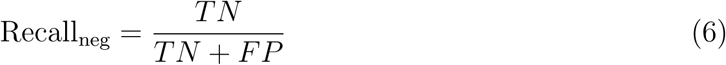

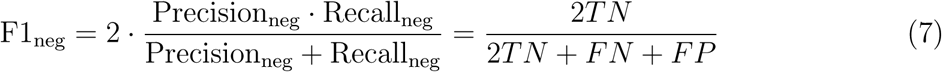

**Matthews Correlation Coefficient (MCC):**

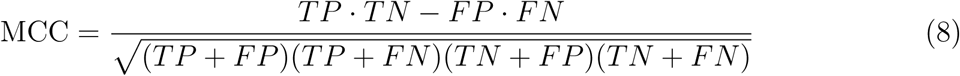

These metrics were computed for all models to quantitatively assess classification accuracy and enable fair comparison between the LoRA-fine-tuned ESM2 and PTMGPT2 model.

### Attention-based Motif Analysis

The optimal fine-tuned ESM2 model was used to extract attention weights by performing forward passes with attention outputs for each protein sequence in the dataset. Sequences were tokenized, and the full attention tensors were obtained in evaluation mode. From these, only the final transformer layer was considered, comprising 40 attention heads. To characterize phosphorylation-associated sequence patterns, attention values were aggregated across all annotated phosphorylation sites. For each attention head independently, a positional window spanning −10 to +10 residues relative to each phosphorylation site was considered. For every position within this window, attention scores from the phosphorylation site to surrounding residues were accumulated conditioned on the amino acid identity, yielding a 21 × 20 matrix representing positional amino acid preferences across all sites. The aggregated attention matrices were converted to probability distributions by row-wise normalization, where each row sum was divided into individual elements to obtain relative frequencies of amino acids at each position. Positions with zero total attention were assigned a uniform distribution (1/20 for each amino acid) as a fallback. The probability matrix was then transformed into an information content matrix using logomaker’s^77^ probability-to-information conversion, which computes bits as log2(probability/background) where background frequencies were assumed uniform (0.05 per amino acid). The information content matrices were visualized as sequence logos^78^ with amino acid letters stacked proportionally to their information content at each position, using a custom color scheme to distinguish the 20 amino acids.

### Evolutionary analysis

For evolutionary analysis, human phosphorylation data were obtained from the EPSD2 database, including protein sequences and annotated phosphorylation sites. Only canonical protein sequences were retained by excluding isoforms, resulting in a curated FASTA dataset of human proteins. To reduce sequence redundancy and group homologous proteins, clustering was performed using CD-HIT^79^ with a sequence identity threshold of 70%. This procedure yielded 20,933 clusters comprising a total of 48,570 sequences, of which 12,451 clusters (59.5%) were singletons. The resulting file was parsed to extract cluster-level statistics, including cluster size, representative sequence, representative sequence length, and mean pairwise identity of non-representative members to the representative. A diversity score for each cluster was defined as 1 − mean identity*/*100. To obtain a high-quality and diverse subset suitable for downstream analysis, clusters were filtered based on the following criteria: (i) cluster size ≥ 3 to ensure sufficient intra-cluster representation, (ii) mean sequence identity *<* 90% to retain sequence diversity, and (iii) representative sequence length between 100 and 1500 amino acids to exclude very short or excessively long proteins. After filtering, 1,369 clusters were retained. The selected clusters exhibited a median size of 4 sequences (mean 6.2, maximum 410) and an average diversity of 0.162. Cluster sizes were predominantly in the range of 3-5 sequences, with a smaller number of larger clusters. For each selected cluster, the representative sequence (as defined by CD-HIT) was extracted and used for subsequent analysis, yielding a non-redundant and evolutionarily diverse set of protein sequences.

The representative sequences from the 1,369 selected clusters were used as queries for large-scale homology search against a diverse set of eukaryotic proteomes. A concatenated protein database comprising proteomes from 200 eukaryotic organisms was constructed by merging individual FASTA files into a single reference file. From this concatenated dataset, organism identifiers were parsed from FASTA headers and a non-redundant list of organisms was generated.

A BLAST^80^ protein database was created from the concatenated proteome. To enable scalable computation, the query FASTA file containing the 1,369 representative human sequences was split into individual single-sequence FASTA files, each corresponding to one query protein. BLASTP searches were performed independently for each query sequence against the concatenated database using an E-value cutoff of 1 × 10*^−^*^5^.^81^ For each query, all potential hits were retrieved, and results were processed to identify the best match per organism. Specifically, for every organism in the dataset, hits were filtered based on organism annotation in the subject title field, and the optimal hit was selected based on the highest bit score, with E-value used as a secondary criterion in the case of ties. This procedure resulted in a per-query mapping of the best homologous sequence across all 200 eukaryotic organisms.

To enable comparison of phosphorylation sites between human proteins and their homologs across 200 eukaryotic organisms, multiple sequence alignments were performed for each set of homologous sequences using the MAFFT^82^ alignment program. For each representative human protein, a corresponding FASTA file containing the homologs identified from BLASTP was used as input. This procedure resulted in a collection of multiple sequence alignments for each human protein and its homologs, enabling positional correspondence analysis of phosphorylation sites across species.

To quantify evolutionary conservation of phosphorylation-related features in the learned representation space, residue-level embeddings were extracted from the fine-tuned model for serine, threonine, and tyrosine residues in both human proteins and their aligned homologous sequences. For each human protein, embeddings corresponding to the aligned phosphorylation sites were identified using the multiple sequence alignments, ensuring positional correspondence between the human sequence and each homolog.

For each aligned residue pair, similarity between the human and organism embeddings was computed using cosine similarity. Given two embedding vectors u and v, the similarity was calculated as

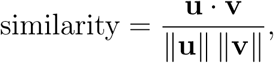

where u · v denotes the dot product and ‖·‖ represents the Euclidean norm. To ensure numerical stability, cases where either vector had zero norm were assigned a similarity score of zero.

For each human protein and corresponding organism, cosine similarity values were computed across all aligned phosphorylation sites and stored. These values were then aggregated at the organism level by pooling similarity scores across all proteins and computing the mean similarity for each organism. This resulted in a single representative similarity score per organism, reflecting the overall conservation of phosphorylation-related embedding features relative to human proteins.

To relate embedding similarity with evolutionary divergence, phylogenetic distances for each organism were obtained from an external dataset. Organism-wise average similarity scores were matched with their corresponding phylogenetic distances and sorted in ascending order of distance from human. The results were visualized using a bar plot, where each bar represents an organism, the height corresponds to the average cosine similarity, and the color encodes phylogenetic distance using a continuous colormap. This analysis enabled systematic assessment of how embedding similarity varies as a function of evolutionary distance.

### Construction of Quantitatively Comparable Embedding and Interaction Graphs

Gene Ontology (GO) data were obtained from the go-basic.obo ontology file, ^83,84^ and human protein annotations were retrieved from the goa_human.gaf dataset.^85^ Only biological process (BP) annotations supported by experimental evidence were retained by excluding entries with the IEA evidence code and those annotated with the NOT qualifier. Protein identifiers were restricted to the set of unique human UniProt accessions present in the EPSD 2.0 database. For each protein, direct GO annotations were expanded by traversing the GO hierarchy using is_a and part_of relationships, thereby incorporating ancestral terms. A mapping from GO terms to associated proteins was constructed, and leaf terms were defined as those without annotated children within the filtered dataset. Only leaf GO terms associated with at least three proteins were retained for subsequent analysis.

Protein-protein interaction data were obtained from STRING v12.0,^86,87^ retaining only interactions with a confidence score of at least 700. STRING identifiers were mapped to UniProt accessions using the provided alias file, and an undirected graph was constructed in which nodes represent proteins and edges correspond to high-confidence interactions. This graph serves as a reference network reflecting experimentally supported functional relationships.

Protein representations were derived from embeddings generated using the fine-tuned ESM2-3B language model. For each protein, residue-level embeddings were averaged to produce a fixed-length vector, and all vectors were L2-normalized such that the cosine similarity between two proteins corresponds to the dot product of their embeddings. To construct an embedding-based similarity network, pairwise cosine similarities were computed and edges were defined by thresholding these similarities. The similarity threshold was calibrated to match the edge density of the STRING network as closely as possible, following principles used in prior network comparison studies. ^64^ Specifically, a random subset of up to 3,000 proteins common to both datasets was sampled with a fixed random seed for reproducibility, and the threshold was chosen as the cosine similarity quantile corresponding to the edge density of the STRING subgraph on those proteins. The calibrated threshold was subsequently constrained to the interval [0.5, 0.99] to ensure numerical validity. The final embedding-derived network was constructed using only proteins shared between the STRING and embedding datasets.

To quantify functional coherence, GO term-specific subgraphs were extracted and analyzed using three graph-based metrics.^88^ The density (*D*) was defined as the ratio of observed edges to the maximum possible number of edges within the induced subgraph. The clustering coefficient (*CC*) was computed as the average local clustering coefficient across nodes within the induced subgraph of each GO term,^89^ considering only edges between proteins annotated to that term. Nodes with degree one within the induced subgraph were assigned a local clustering coefficient of one, as their single neighbor trivially satisfies the clustering condition, while isolated nodes were assigned a value of zero. Functional segregation was quantified using the segregation ratio (*S_r_*), defined as

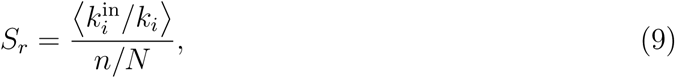

where 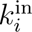 denotes the number of neighbors of node *i* within the GO term subgraph, *k_i_* is its total degree in the full graph, *n* is the size of the GO term, and *N* is the total number of nodes. Proteins with zero degree in the full graph were excluded from the segregation ratio computation.

Leaf GO terms were grouped into higher-level biological process categories based on the GO hierarchy, and all three metrics were averaged across leaf terms within each category. The number of leaf terms, average term size, and mean values of *S_r_*, *CC*, and *D* were reported for both the STRING network and the embedding-derived network, enabling a direct and controlled comparison of their ability to capture biologically meaningful functional organization.

## Code and Data Availability

The code for PTM site prediction models is publicly available at https://github.com/JMLab-tifrh/PTM_prediction. The repository includes pre-trained models for acetylation, phosphorylation, and ubiquitination site prediction, along with Jupyter notebooks for running predictions on new protein sequences. The datasets used for model training and evaluation were obtained from publicly available databases: EPSD 2.0 https://epsd.biocuckoo. cn/Download.php#1 and dbPTM https://biomics.lab.nycu.edu.tw/dbPTM/download.php#dataset.

## Supporting Information

The Supporting Information (SI) provides supplemental figures, supplemental table. Figure S1 shows the comparison of base ESM2 models with PTMGPT2 model. Figure S2 shows the Logo map for kinase specific motifs. Figure S3 and S4 shows the cosine similarity of threonine and tyrosine phosphorylation across eukaryotic species. Table S1 shows the comparison of STRING and ESM2 network derived metrics for different GO terms.

## Acknowledgement

We acknowledge support of the Department of Atomic Energy, Government of India, under Project Identification No. RTI 4007. We sincerely acknowledge Tata Institute of Fundamental Research Hyderabad, India for providing the support of computing resources. JM acknowledges the core research grant approved by the Department of Science and Technology (DST) of India (CRG/2023/001426).

