## Supplemental Figures for "Task-Specialized Protein Language Models Decode the Sequence Grammar of Post-Translational Modification Sites"

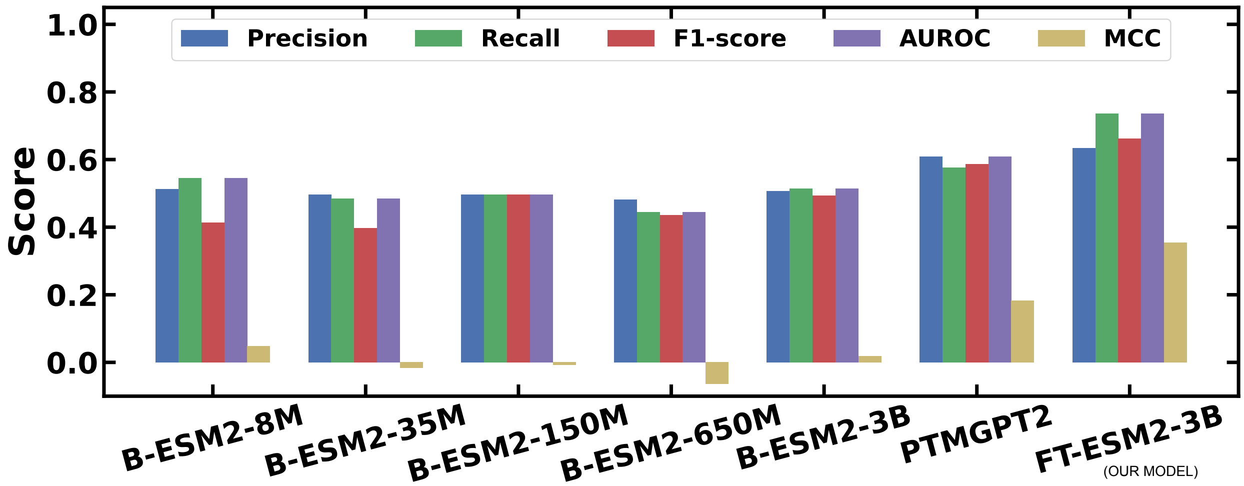


**Figure S1:** Comparison of the classification metrics for different base ESM2 models (i.e. without finetuning) (referred as *B-ESM2*) for phosphorylation and compared with PTMGPT2 model and this work's fine-tuned ESM2-3B model (*FT-ESM2-3B*)

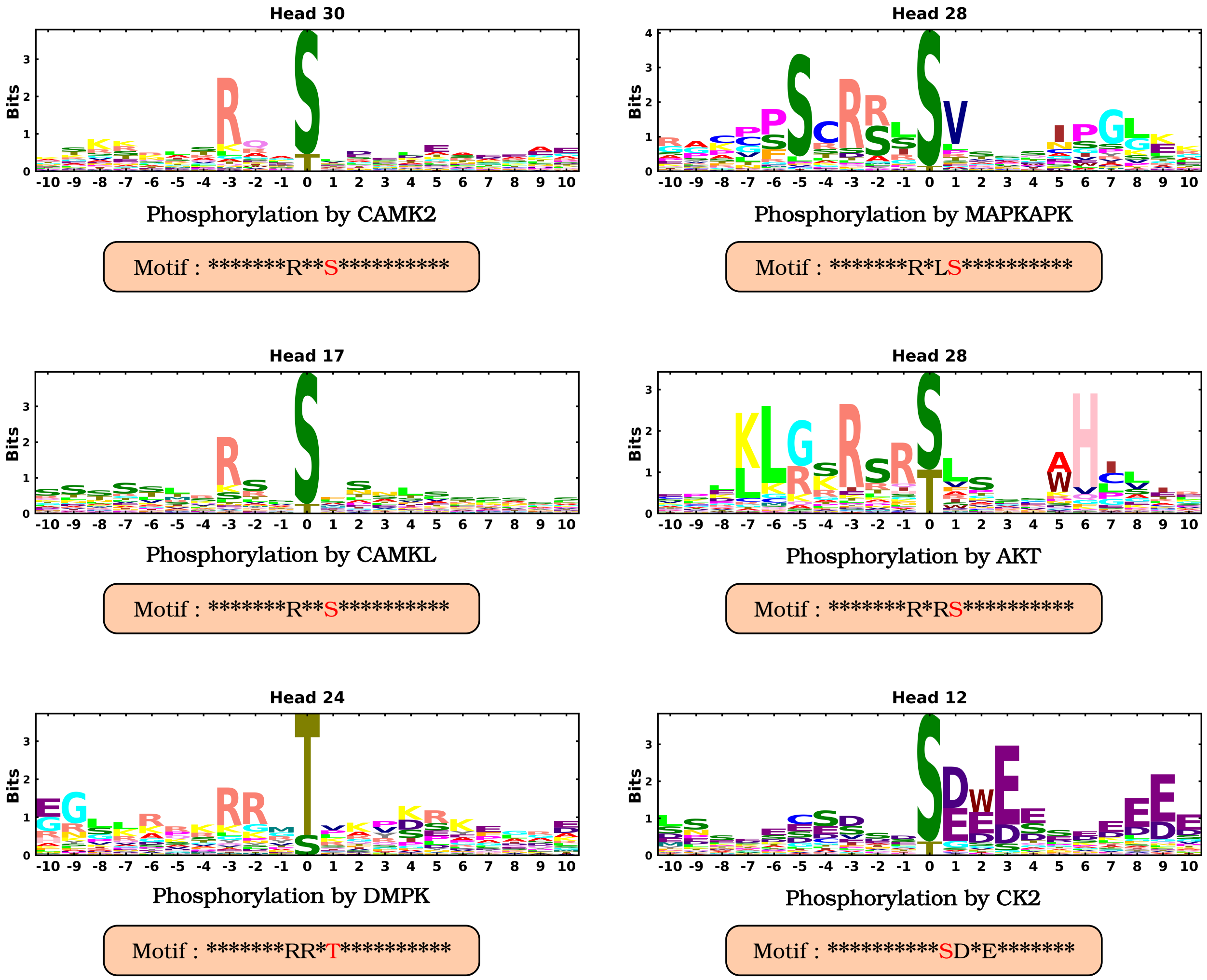


**Figure S2:** Sequence logos derived from attention-based information content highlight enriched residue patterns surrounding phosphorylation sites, revealing kinase-specific motifs (e.g., CAMK2, CAMKL, DMPK, MAPKAPK, AKT, CK2) captured by the model.

**Table S1:** Comparison between STRING and ESM2 networks

| ID | Name | Np | Size | Sr_STRING | CC_STRING | D_STRING | Np_ESM | Size_ESM | Sr_ESM | CC_ESM | D_ESM | ΔSr | ΔCC | ΔD |
|---|---|---|---|---|---|---|---|---|---|---|---|---|---|---|
| GO:0051301 | cell division | 9 | 14.70 | 484.41 | 0.55 | 0.40 | 9 | 14.70 | 775.06 | 0.39 | 0.23 | 290.65 | -0.16 | -0.17 |
| GO:0051234 | establishment of localization | 255 | 9.20 | 345.53 | 0.53 | 0.31 | 255 | 9.20 | 869.71 | 0.33 | 0.13 | 524.18 | -0.20 | -0.18 |
| GO:0008152 | metabolic process | 536 | 11.40 | 328.61 | 0.67 | 0.46 | 536 | 11.40 | 693.94 | 0.22 | 0.08 | 365.33 | -0.45 | -0.38 |
| GO:0051179 | localization | 308 | 9.60 | 322.69 | 0.53 | 0.32 | 308 | 9.60 | 796.67 | 0.31 | 0.12 | 473.98 | -0.22 | -0.20 |
| GO:0007049 | cell cycle | 63 | 23.70 | 295.02 | 0.64 | 0.37 | 63 | 23.70 | 267.54 | 0.21 | 0.06 | -27.48 | -0.43 | -0.31 |
| GO:0009987 | cellular process | 1578 | 12.60 | 251.69 | 0.56 | 0.35 | 1578 | 12.60 | 506.84 | 0.24 | 0.08 | 255.15 | -0.32 | -0.27 |
| GO:0016043 | cellular component organization | 354 | 13.90 | 242.98 | 0.57 | 0.34 | 354 | 13.90 | 372.47 | 0.23 | 0.07 | 129.49 | -0.34 | -0.27 |
| GO:0007154 | cell communication | 234 | 11.80 | 224.02 | 0.64 | 0.41 | 234 | 11.80 | 339.47 | 0.28 | 0.08 | 115.45 | -0.36 | -0.33 |
| GO:0022414 | reproductive process | 80 | 38.50 | 215.84 | 0.43 | 0.20 | 80 | 38.50 | 228.03 | 0.19 | 0.04 | 12.19 | -0.24 | -0.16 |
| GO:0019725 | cellular homeostasis | 29 | 11.40 | 192.21 | 0.47 | 0.24 | 29 | 11.40 | 517.51 | 0.18 | 0.05 | 325.30 | -0.29 | -0.19 |
| GO:0050896 | response to stimulus | 481 | 14.60 | 182.81 | 0.57 | 0.31 | 481 | 14.60 | 356.40 | 0.24 | 0.07 | 173.59 | -0.33 | -0.24 |
| GO:0040007 | growth | 19 | 5.00 | 134.75 | 0.27 | 0.14 | 19 | 5.00 | 443.07 | 0.16 | 0.05 | 308.32 | -0.11 | -0.09 |
| GO:0065007 | biological regulation | 1224 | 11.20 | 130.37 | 0.45 | 0.24 | 1224 | 11.20 | 263.60 | 0.18 | 0.05 | 133.23 | -0.27 | -0.19 |
| GO:0002376 | immune system process | 116 | 13.90 | 117.87 | 0.53 | 0.27 | 116 | 13.90 | 369.08 | 0.23 | 0.07 | 251.21 | -0.30 | -0.20 |
| GO:0048518 | positive regulation of biological process | 515 | 11.70 | 105.21 | 0.43 | 0.20 | 515 | 11.70 | 238.55 | 0.16 | 0.04 | 133.34 | -0.27 | -0.16 |
| GO:0048519 | negative regulation of biological process | 382 | 9.50 | 98.11 | 0.36 | 0.17 | 382 | 9.50 | 225.48 | 0.13 | 0.04 | 127.37 | -0.23 | -0.13 |
| GO:0032502 | developmental process | 599 | 20.30 | 91.17 | 0.36 | 0.13 | 599 | 20.30 | 144.33 | 0.21 | 0.06 | 53.16 | -0.15 | -0.07 |

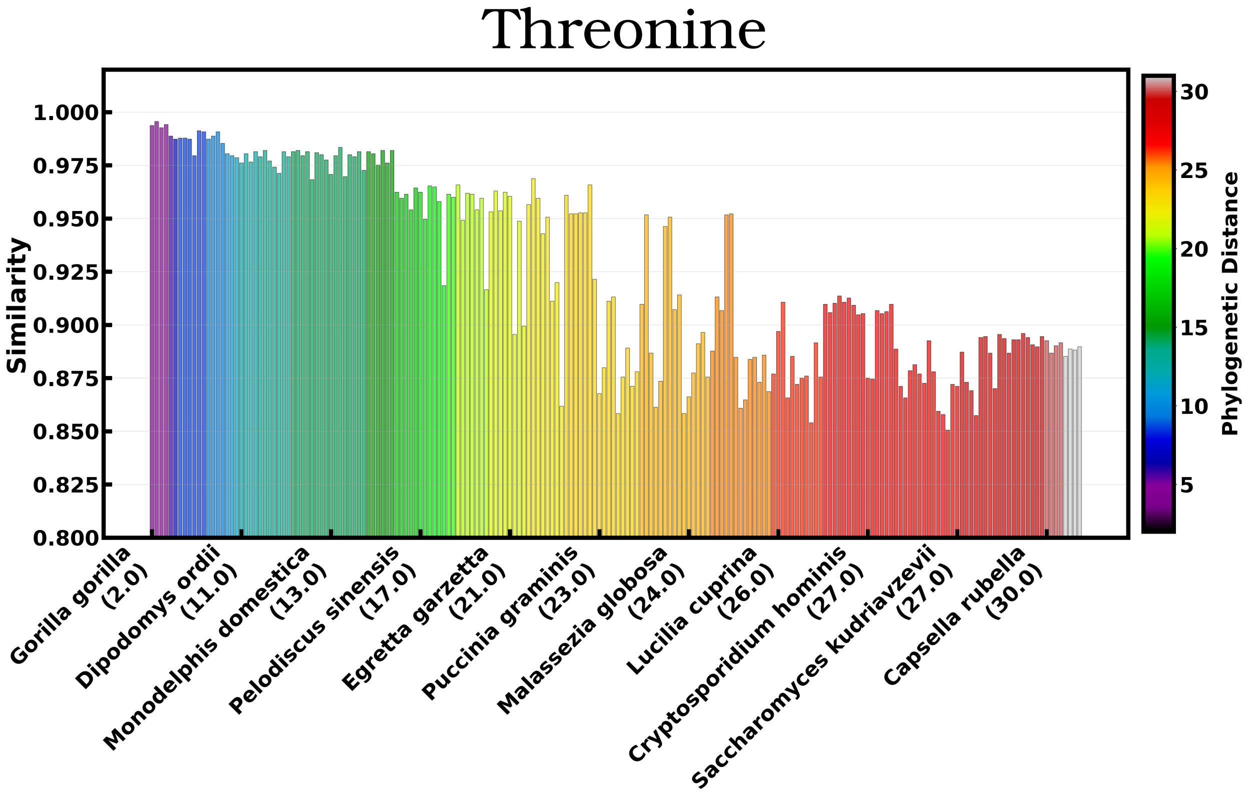


**Figure S3:** Cosine similarity of threonine phosphorylation site embeddings across diverse eukaryotic species, organized according to phylogenetic distance to reflect evolutionary divergence patterns.

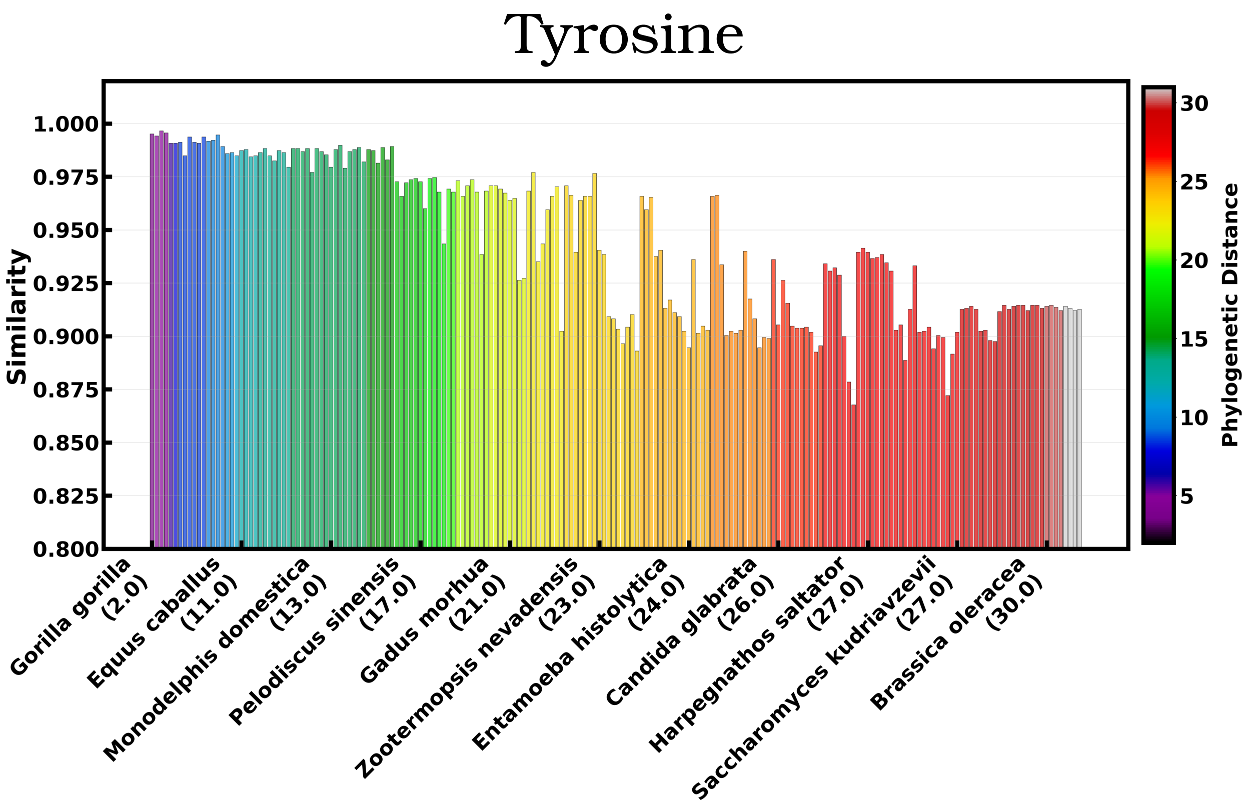


**Figure S4:** Cosine similarity of tyrosine phosphorylation site embeddings across diverse eukaryotic species, organized according to phylogenetic distance to reflect evolutionary divergence patterns.
